# Contact inhibitory Eph signaling suppresses EGF-promoted cell migration by decoupling EGFR activity from vesicular recycling

**DOI:** 10.1101/202705

**Authors:** Wayne Stallaert, Ola Sabet, Yannick Brüggemann, Lisa Baak, Philippe I.H. Bastiaens

## Abstract

The ability of cells to adapt their behavior to growth factors in relation to their environment is an essential aspect of tissue development and homeostasis. Here we show that Eph receptor signaling from cell-cell contacts changes the cellular response to EGFR activation by altering its vesicular trafficking. Eph receptor activation traps EGFR in Rab5-positive early endosomes through an inhibition of Akt-dependent vesicular recycling. By altering the spatial distribution of EGFR activity during EGF stimulation, Eph receptor activation selectively suppresses migratory Akt signaling from the plasma membrane, while preserving proliferative ERK signaling from endosomes. We also show that soluble extracellular signals engaging the G-protein coupled receptor Kiss1 similarly suppress vesicular recycling to alter EGFR signaling. The cellular environment can thus modulate EGFR vesicular trafficking dynamics to generate context-dependent responses to EGF stimulation.

**Summary:** Eph receptor activation generates context-dependent cellular responses to EGFR activation by altering its vesicular trafficking dynamics.

## Introduction

Activation of epidermal growth factor receptor (EGFR) promotes a variety of cellular responses including cell growth, proliferation, survival, apoptosis, differentiation and migration (*1*), some of which are functionally opposed. To select among these diverse outcomes, the cell requires additional contextual information. This context can be intrinsic (e.g. cell type or cell cycle stage), or extrinsic, in the form of extracellular signals that provide information about the current (or past) environmental context. Adaptability to a changing environment requires that extrinsic information be integrated through mechanisms that can transform the response to subsequent growth factor stimulation.

Local cell density is one such example of extrinsic context that can influence cellular activity to generate distinct functional states (*2*-*4*). The Eph family of receptor tyrosine kinases act as sensors of cell density, becoming activated at points of cell-cell contact through interactions with membrane bound ephrin ligands presented on the surfaces of adjacent cells (*5*). Eph receptors in many ways operate in functional opposition to EGFR, acting as tumour suppressors (*5*-*11*) and mediating contact inhibition of locomotion to suppress cellular migration and metastasis (*12*-*15*). Moreover, a functional coupling of EGFR and Eph receptor activity has been shown to control cell migration (*15*). Although the precise mechanism through which Eph receptors regulate EGF-promoted migration remains elusive, a convergence of receptor activity on phosphoinositide 3-kinase (PI3K)/Akt signaling was implicated.

Akt has been shown to regulate EGFR vesicular trafficking through the endosomal system (*16*). By stimulating the activity of the early endosomal effector PIKfyve (FYVE-containing phosphatidylinositol 3-phosphate 5-kinase), Akt activity controls the transition of EGFR through early endosomes, regulating both its recycling back to the PM and its degradation in the lysosome. Thus, while endocytosis of cell surface receptors has traditionally been viewed as a mechanism to attenuate downstream signaling following ligand stimulation, the notion that signaling molecules downstream of cell surface receptors can, in turn, influence vesicular trafficking (*16*-*21*) generates a reciprocal relationship between receptor activation and vesicular dynamics whose role in shaping the cellular response to stimuli has only recently begun to garner attention (*22*). Furthermore, this bidirectional relationship could also allow the signaling activity of one receptor to influence the response properties of another through changes in its vesicular trafficking dynamics, generating context-dependent receptor activity.

In the current work, we show that Eph receptor activation at cell-cell contacts regulates the vesicular dynamics of EGFR through an inhibition of Akt-dependent trafficking. By modulating the spatial distribution of EGFR activity, Eph receptor activation alters the cellular response to EGF stimulation, selectively suppressing EGF-promoted migratory signaling while preserving its effect on proliferation.

## RESULTS

### Eph receptor activation affects EGFR vesicular trafficking

Stimulation of endogenous Eph receptors in Cos-7 cells with a soluble, clustered ephrinA1-Fc (A1) ligand (*23*, *24*) induced a reduction in EGFR abundance at the PM (**Fig. 1A-B**). This decrease in PM EGFR abundance following Eph receptor activation was observed in various cell lines, including HEK293, NIH 3T3, MCF10A and MDA-MB-231 cells (**Fig. S1A-D**), exhibiting a wide range of endogenous EGFR expression (**Fig. S1E**).

**Figure 1:**
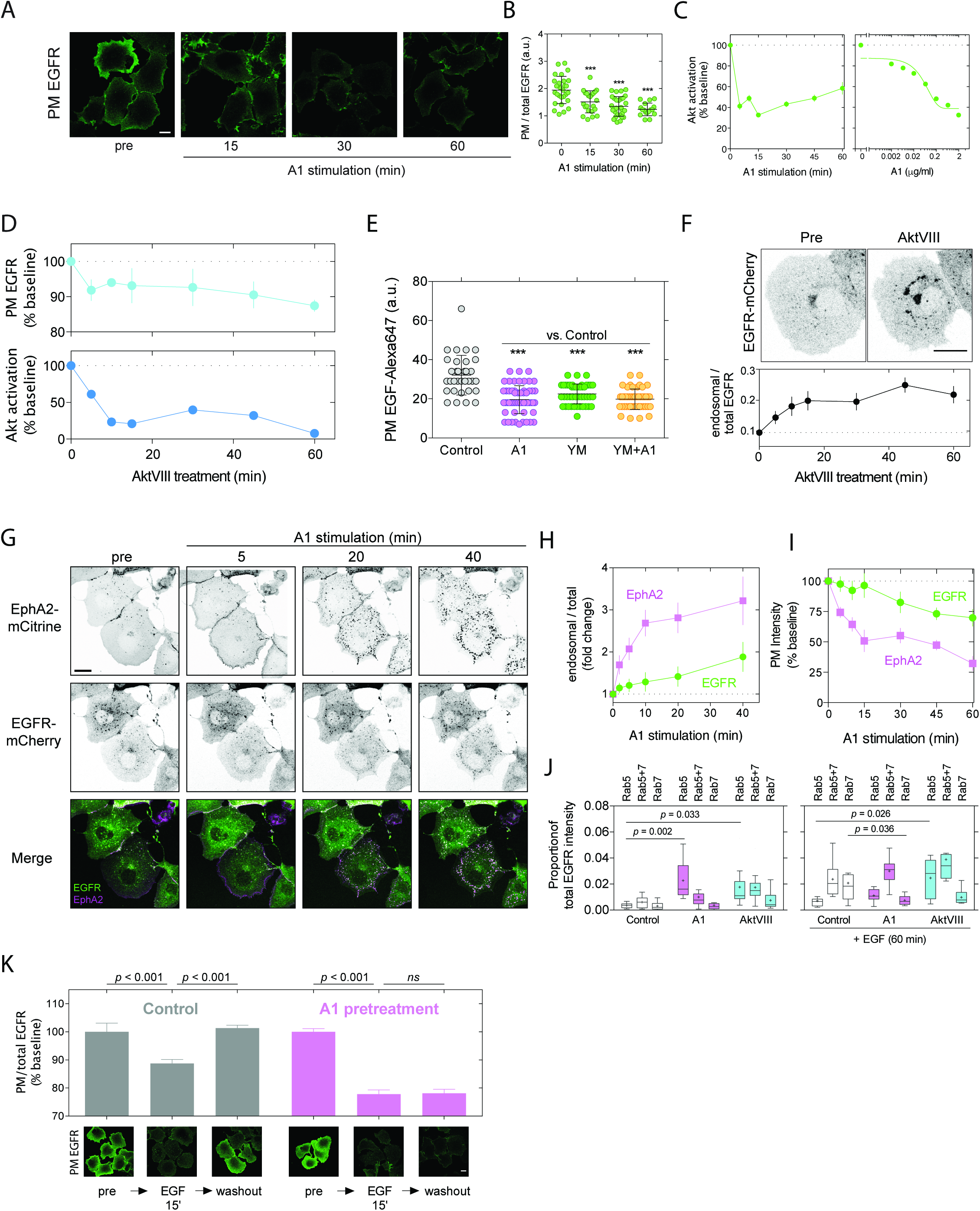
Eph receptor activation affects Akt-dependent EGFR trafficking. **(A,B)** Representative immunofluorescence images (A) and quantification (B) of endogenous PM EGFR in Cos-7 cells (*N* = 16-29 cells/condition) following stimulation with ephrinA1-Fc (A1, 2 μg/ml) for the indicated times (means ± sd). **(C)** Quantification of Akt activation by In-cell Western (ICW) in Cos-7 cells following A1 stimulation (left: 2 μg/ml; right: 15 min) (means ± s.e.m.). **(D)** Quantification of endogenous PM EGFR abundance by On-cell Western (OCW) and Akt activation in Cos-7 cells following treatment with the Akt inhibitor AktVIII (10 μM) for the times indicated (means ± s.e.m.). **(E)** EGF-Alexa647 binding (200 ng/ml, 2 min) to endogenous EGFR in Cos-7 cells as a measure of PM EGFR abundance following 1 h pretreatment with A1 (2 μg/ml), the PIKfyve inhibitor YM201636 (YM, 200 nM) or both (*N*=33-59 cells/condition) (means ± sd). **(F)** Representative confocal images of Cos-7 cells expressing EGFR-mCherry before (top left) and after (top right) treatment with AktVIII (10 μM, 1 h) and quantification of the increase in endosomal EGFR-mCherry during AktVIII treatment (bottom, *N* = 6 cells, means ± sd). **(G)** Representative time-lapse confocal images of Cos-7 cells expressing EGFR-mCherry and EphA2-mCitrine following A1 stimulation (2 μg/ml). **(H)** Quantification of endosomal EGFR-mCherry and EphA2-mCitrine from time-lapse confocal imaging (G) during A1 stimulation (*N* = 7 cells, means ± sd). **(I)** Quantification of PM EGFR-mCherry and EphA2-mCitrine abundance by OCW during A1 stimulation (2 μg/ml) (means ± s.e.m.). **(J)** Immunofluorescence measurements of EGFR intensity in Rab5-, Rab5/Rab7-and Rab7-positive endosomal compartments in control, A1-(2 μg/ml, 1 h) and AktVIII-(10 μM, 1 h) pretreated Cos-7 cells prior to (left) and following EGF stimulation (right, 100 ng/ml, 1 h). (*N*=6-11 cells/condition). Data are represented by Tukey boxplots with the mean denoted as a cross and the median as a line. **(K)** Measurements of EGFR recycling in control and A1-pretreated cells (2 μg/ml, 1 h) by immunofluorescence prior to (pre), after EGF stimulation (10 ng/ml, 15 min) and 15 min following EGF washout (*N* = 34-40 cells/condition, means ± sd). Statistical significance was determined in B, E, J and K using a one-way ANOVA with Sidak’s *post-hoc* test (***, *p <* 0.001). Scale bars = 20 μm.

While EGFR internalization is a well-established consequence of growth factor-induced receptor activation, we observed that the loss of PM EGFR did not result from an Eph receptor-induced transactivation of EGFR (**Fig S1F**). Since EGFR also continuously recycles through the endosomal system in the absence of growth factor stimulation (*25*), we hypothesized that Eph receptor activation may reduce PM EGFR abundance by trapping constitutively recycling receptors in endosomes.

Eph receptor activation decreases the activity of Akt (**Fig. *1C***) (*14*), a signaling effector previously demonstrated to promote EGFR vesicular recycling (*16*). Direct inhibition of Akt or its downstream early endosomal effector PIKfyve (*16*) indeed reduced PM EGFR abundance in Cos-7 cells (**Fig. 1D-E**). A1 stimulation or PIKfyve inhibition promoted similar decreases in PM EGFR abundance, as measured by a reduction in PM EGF-Alexa647 binding (**Fig. 1E**). Furthermore, the combination of Eph receptor activation with PIKfyve inhibition did not further reduce PM EGFR abundance (**Fig. 1E**), indicative of a shared molecular mechanism. Consistent with a suppression of constitutive EGFR recycling, we observed an endosomal accumulation of ectopically expressed EGFR-mCherry in live cells after Akt or PIKfyve inhibition (**Fig. 1F**, **Movie S1**, **Fig. S2A-B**). Time lapse confocal imaging of Cos-7 cells expressing EGFR-mCherry and EphA2-mCitrine also revealed an endosomal accumulation of EGFR with time following soluble A1 stimulation (**Fig. 1G-I**, **Movie S2**) or upon presentation of ephrinA1 ligand on the membrane of adjacent cells at sites of cell-cell contact (**Movie S3**). This shift in the spatial distribution of EGFR from the PM to endosomes following A1 stimulation occurred primarily by trapping receptors in Rab5-positive early endosomes (**Fig. 1J**), consistent with an inhibition of Akt-dependent trafficking (*16*) (**Fig. 1J**). Thus, Eph receptor activation alters the subcellular distribution of EGFR prior to growth factor stimulation by trapping constitutively recycling receptors in Rab5-positive early endosomes through an inhibition of Akt/PIKfyve-dependent vesicular recycling.

We next investigated how Eph receptor activation influences EGFR trafficking during EGF stimulation. The trafficking fate of EGFR through the endosomal system is determined by post-translational modifications, with receptor ubiquitination acting as a molecular switch that diverts EGFR to the lysosome for degradation (*25*). Since EGFR ubiquitination increases with EGF binding (*26*), saturating EGF concentrations (> 50 ng/ml (*27*)) generate a finite temporal signaling response by progressively depleting PM EGFR through ubiquitin-dependent lysosomal degradation. Stimulation of endogenous receptors in Cos-7 cells with a saturating concentration of EGF (100 ng/ml) induced a ~40% reduction in total EGFR expression after 60 min of stimulation (**Fig. S2C**) and residual EGFR resided primarily in Rab7-positive late endosomes (**Fig. 1J**). A1 pretreatment or direct Akt inhibition, in contrast, impaired Rab5-to-Rab7 endosomal maturation (*28*) (**Fig. 1J**), leading to a reduction in receptor degradation at saturating EGF concentrations (≥50 ng/ml; **Fig. S2C**).

At lower, subsaturating EGF concentrations typically found in human tissue secretions (0.4-20 ng/ml)(*29*), only a fraction of receptors are ligand bound, receptor ubiquitination is reduced (*26*), and internalized receptors are preferentially recycled back to the PM (*30*). To assess whether Eph receptors inhibit EGFR recycling following subsaturating EGF stimulation, we pulsed endogenous receptors in Cos-7 cells with 10 ng/ml EGF to induce EGFR endocytosis and measured its subsequent return to the PM following EGF washout (**Fig. 1K**). While we observed a complete recovery of PM EGFR abundance in control cells following EGF washout, A1 pretreatment completely suppressed EGFR recycling.

Thus, Eph receptor activation suppresses EGFR trafficking through the early endosome during EGF stimulation; impairing the recycling of non-ubiquitinated receptors back to the PM as well as inhibiting the transition of ubiquitinated receptors to late endosomes.

### Eph receptor activation changes the spatial distribution of EGFR activity

Many functional outcomes to EGFR activation, such as cellular migration, require that cells remain responsive to persistent growth factor stimulation. To ensure sensitivity to stimuli during long periods of exposure, the cell must maintain sufficient receptor abundance at the PM despite the continuous internalization of activated receptors. We therefore posed the following questions: Does Akt-dependent recycling help maintain cellular responsiveness to EGF during persistent, subsaturating stimulation? Can Eph receptor activation at cell-cell contacts change the response properties of EGFR by modulating its vesicular trafficking?

To address the impact of Akt-dependent recycling on EGFR activation, measurements of endogenous EGFR phosphorylation and trafficking in Cos-7 cells were obtained by immunofluorescence following subsaturating EGF stimulation in control cells and following Eph receptor activation or Akt/PIKfyve inhibition. Individual cells were radially segmented to quantify changes in the average spatial distribution of EGFR activity with time and visualized using 3-D spatial-temporal maps (**Fig. 2A**). Through an accumulation of EGFR in endosomal compartments during sustained EGF stimulation, cells pretreated with either A1 or an Akt inhibitor generated less EGFR phosphorylation after 60 min of EGF stimulation relative to control cells (**Fig. 2A-B**). Decoupling Akt activation from its effect on trafficking by PIKfyve inhibition had indistinguishable effects from direct Akt inhibition or A1 pretreatment on EGFR phosphorylation and trafficking (**Fig. 2A-B**), indicating that Akt activity maintains EGFR activation at the PM during sustained, subsaturating EGF stimulation through its effects on vesicular recycling.

**Figure 2:**
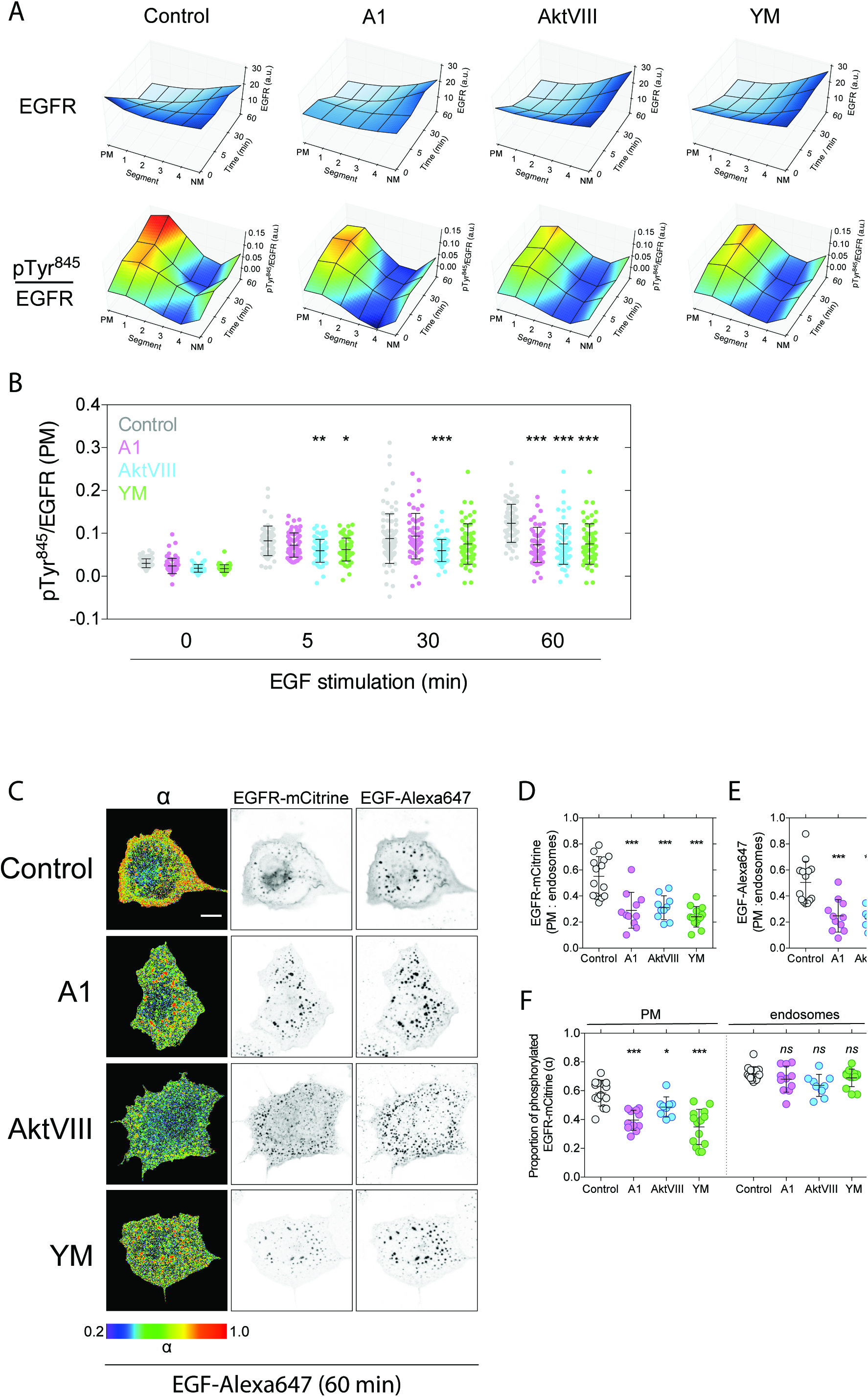
Eph receptor activation changes the spatial distribution of EGFR activity. **(A-B)** Average spatial-temporal maps (A) of endogenous EGFR abundance (top) and Tyr845 phosphorylation (bottom) in radially segmented Cos-7 cells (plasma membrane, PM **->** nuclear membrane, NM) prior to and during EGF stimulation (20 ng/ml for 5, 30 and 60 min) in control, ephrinA1-Fc-(A1, 2 μg/ml, 1 h), AktVIII-(10 μM, 1 h) and YM201636-(YM, 200 nM, 1h) pretreated cells (*N* = 50-90 cells per condition). Single cell measurements of EGFR pTyr^845^ phosphorylation in the PM segment (B) during EGF stimulation (means ± sd) **(C)** Phosphorylated fraction of EGFR-mCitrine as detected by FLIM-FRET (α) and representative images of EGFR-mCitrine and EGF-Alexa647 fluorescence in control and A1-(2 μg/ml, 1 h), AktVIII-(10 μM, 1 h) and YM-(200 nM, 1h) pretreated Cos-7 cells following 60 min of EGF-Alexa647 stimulation (20 ng/ml) (*N* = 10-14 cells/condition). **(D-F)** Quantification of the PM:endosome ratio of (D) EGFR-mCitrine and (E) EGF-Alexa647 fluorescence intensity and **(F)** phosphorylated fraction of EGFR-mCitrine (α) at PM and endosomes (means ± sd). Statistical significance was determined in B and D-F using a one-way ANOVA with Sidak’s *post-hoc* test (***, *p <* 0.001; **, *p <* 0.01; *, *p <* 0.05). Scale bars = 20 μm.

To specifically quantify EGFR phosphorylation at the PM and on endosomes during sustained, subsaturating EGF stimulation, we employed fluorescence lifetime imaging microscopy (FLIM) to detect Förster resonance energy transfer (FRET) between EGFR-mCitrine and a phospho-tyrosine binding domain fused to mCherry (PTB-mCherry) (*31*) in Cos-7 cells (**Fig. 2C-F**). In control cells, EGFR-mCitrine remained highly phosphorylated at both the PM and in endosomes following 60 min of sustained EGF-Alexa647 stimulation. In cells pretreated with A1 or following Akt or PIKfyve inhibition, we observed reduced PM EGFR-mCitrine density (**Fig. 2D**) and EGF-Alexa647 binding (**Fig. 2E**), resulting in diminished EGFR-mCitrine phosphorylation specifically at the PM (**Fig. 2F**). In these conditions in which Akt-dependent recycling was suppressed, ligand-bound, active EGFR-mCitrine accumulated in endosomes, maintaining its phosphorylation in this compartment to the same extent as control cells (**Fig. 2C-F**).

Eph receptor activation, therefore, by inhibiting Akt-dependent recycling, changes the spatial distribution of EGFR activity during sustained, subsaturating EGF stimulation, selectively reducing EGFR activation at the PM while preserving receptor activity in endosomes.

### Eph receptor activation at cell-cell contact alters the EGFR signaling response

Although EGFR continues to activate signaling effectors from endosomal membranes (*32*-*40*), Akt is preferentially activated at the PM (*41*, *42*) (**Fig. S3**). We therefore investigated how Eph receptor activation, by changing the spatial distribution of EGFR activity, regulates its signaling response during EGF stimulation. By suppressing vesicular recycling and reducing EGFR activity at the PM, A1 pretreatment selectively inhibited Akt activation following subsaturating EGF stimulation of endogenous receptors in Cos-7 (**Fig. 3A** **top**, **Fig. S4A**) and HEK293 cells (**Fig. S4B**), while ERK activation, which can continue from endosomal membranes (*37*, *39*, *43*) (**Fig. S3**), remained intact (**Fig 3A** **bottom**, **Fig. S4**). To confirm that EphA2 inhibits EGF-promoted Akt activation by suppressing EGFR recycling and does not simply reflect the opposed regulation of Akt by EGFR and EphA2 (activation vs inhibition, respectively), we assessed whether EGFR trafficking was dispensable for the A1-induced suppression of EGF-promoted Akt activation. Cells were prestimulated with A1, followed by treatment with the dynamin inhibitor dynole 34-2 to block subsequent endocytosis, and then stimulated with EGF. When EGFR endocytosis was blocked (**Fig. S3C**), A1 pretreatment had no effect on EGF-promoted Akt activation (**Fig. 3B** **top**). Pretreatment with the negative control analogue dynole 31-2, to control for off-target effects, did not inhibit EGFR endocytosis (**Fig. S3C**), and had no effect on A1-induced suppression of EGF-promoted Akt activation (**Fig. 3B** **bottom**), corroborating that intact EGFR vesicular trafficking is required for the inhibitory effect of Eph receptors on EGFR signaling.

**Figure 3:**
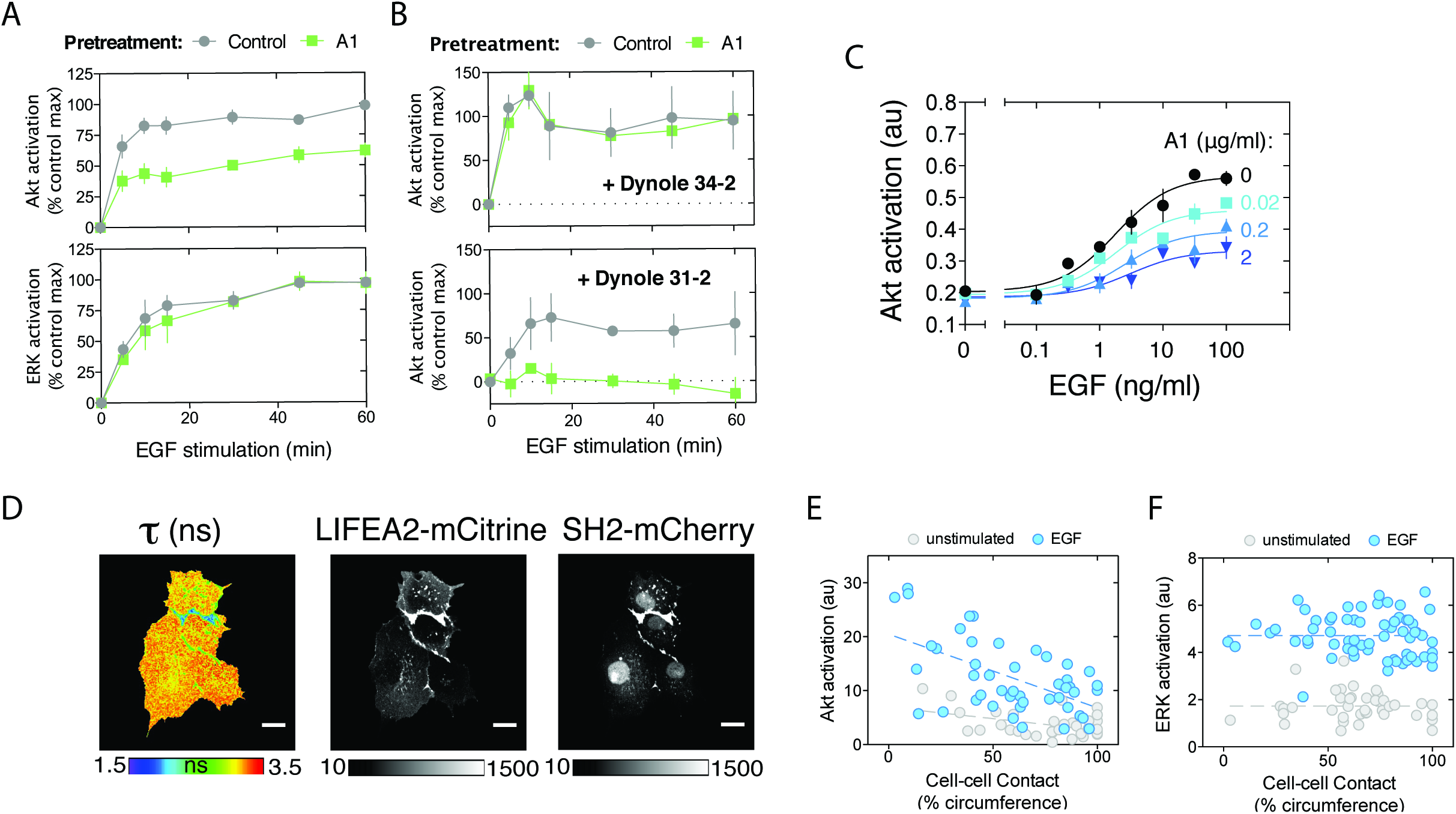
Eph receptor activation at cell-cell contact alters the EGFR signaling response. **(A)** Quantification of Akt (top) and ERK (bottom) activation by ICW in control and ephrinA1-Fc-pretreated (A1, 2 μg/ml, 1 h) Cos-7 cells endogenously expressing the receptors following EGF stimulation (1 ng/ml) (means ± s.e.m.) **(B)** Quantification of Akt activation in control or A1-pretreated (2 μg/ml, 1 h) HEK293 cells, followed by 30 min treatment with the dynamin inhibitor dynole 34-2 (100 μM, top) or its negative control analog dynole 31-2 (100 μM, bottom), then stimulated with EGF (1 ng/ml) for the times indicated (means ± s.e.m.) **(C)** Quantification of EGF-promoted Akt activation in Cos-7 cells following pretreatment with increasing concentrations of A1 (0.02, 0.2 and 2 μg/ml, 1 h) (means ± s.e.m.) **(D)** Representative images of EphA2 activity using a FRET-based sensor (LIFEA2(*24*)), whereby a decrease in fluorescence lifetime (τ, ns) represents an increase in activity, and fluorescence intensity measurements of LIFEA2-mCitrine and SH2-mCherry in Cos-7 cells. Scale bar = 20 μm. **(E,F)** Single cell measurements of Akt (E) and ERK (F) activation versus cell-cell contact in 2-D cultures (% cell circumference) in unstimulated and EGF stimulated (20 ng/ml, 1 h) Cos-7 cells. A sum-of-squares F test was used to determine significance: Akt, unstimulated: *F* = 16.0, *p* = 0.001, r^2^ = 0.432; Akt, EGF: *F=* 21.4, *p <* 0.001, r^2^ = 0.322; ERK, unstimulated: *F* = 0.180, *p* = 0.673, r^2^ = 0.003; ERK, EGF: *F* = 0.321, *p* = 0.575, r^2^ = 0.009.

Increasing concentrations of A1 progressively inhibited EGF-mediated Akt activation (**Fig. 3C**), suggesting that the degree of cell-cell contact might determine the magnitude of Akt activation in response to a given concentration of EGF. To directly investigate the influence of cell-cell contact on EGFR signaling, we obtained single cell measurements of Akt and ERK activation in Cos-7 with varying degrees of cell-cell contact. Homotypic cell-cell contact promotes Eph receptor activation through interactions with ephrins presented on neighboring cells (*24*) (**Fig. 3D**). Akt activation decreased with cell-cell contact both prior to and following EGF stimulation (**Fig. 3E**), demonstrating that increasing cell-cell contact reduces the magnitude of EGF-promoted Akt activation. ERK activation, on the other hand, was unaffected by cell-cell contact, with cells generating similar EGF-promoted increases in ERK activation irrespective of their degree of cell-cell contact (**Fig. 3F**).

### Coupling EGFR activity to vesicular recycling generates a positive feedback

While the inhibition of Akt-dependent recycling results in a reduction in PM EGFR abundance in Cos-7 cells (**Fig. 1A-K**, **Fig. S1A-D**, **Fig. S2A-B**), we also observed that an increase in cellular Akt activity through the inhibition of its negative regulator PP2A by okadaic acid resulted in a concomitant increase in PM EGFR (**Fig. 4A**). Since EGFR activation itself increases Akt activity in cells (**Fig. 3A-C**, **Fig. 3E**, **Fig. S3**), we next asked whether PM EGFR abundance is actively maintained during growth factor stimulation through an EGF-induced increase in Akt-dependent vesicular recycling. Using a fluorescence localization after photoactivation (FLAP) approach to quantify the vesicular trafficking of EGFR to the PM following photoactivation of EGFR-paGFP in endosomes (**Fig. 4B** **top**), we observed an increase in EGFR-paGFP recycling during EGF stimulation (**Fig. 4B** **bottom**). Akt inhibition completely suppressed this EGF-promoted increase in vesicular recycling (**Fig. 4B** **bottom**), further demonstrating the contribution of Akt-dependent recycling in sustaining PM EGFR activity (**Fig. 3A**). Thus, by stimulating Akt-dependent recycling, EGFR activation generates a positive feedback that actively maintain its PM abundance during EGF stimulation.

**Figure 4:**
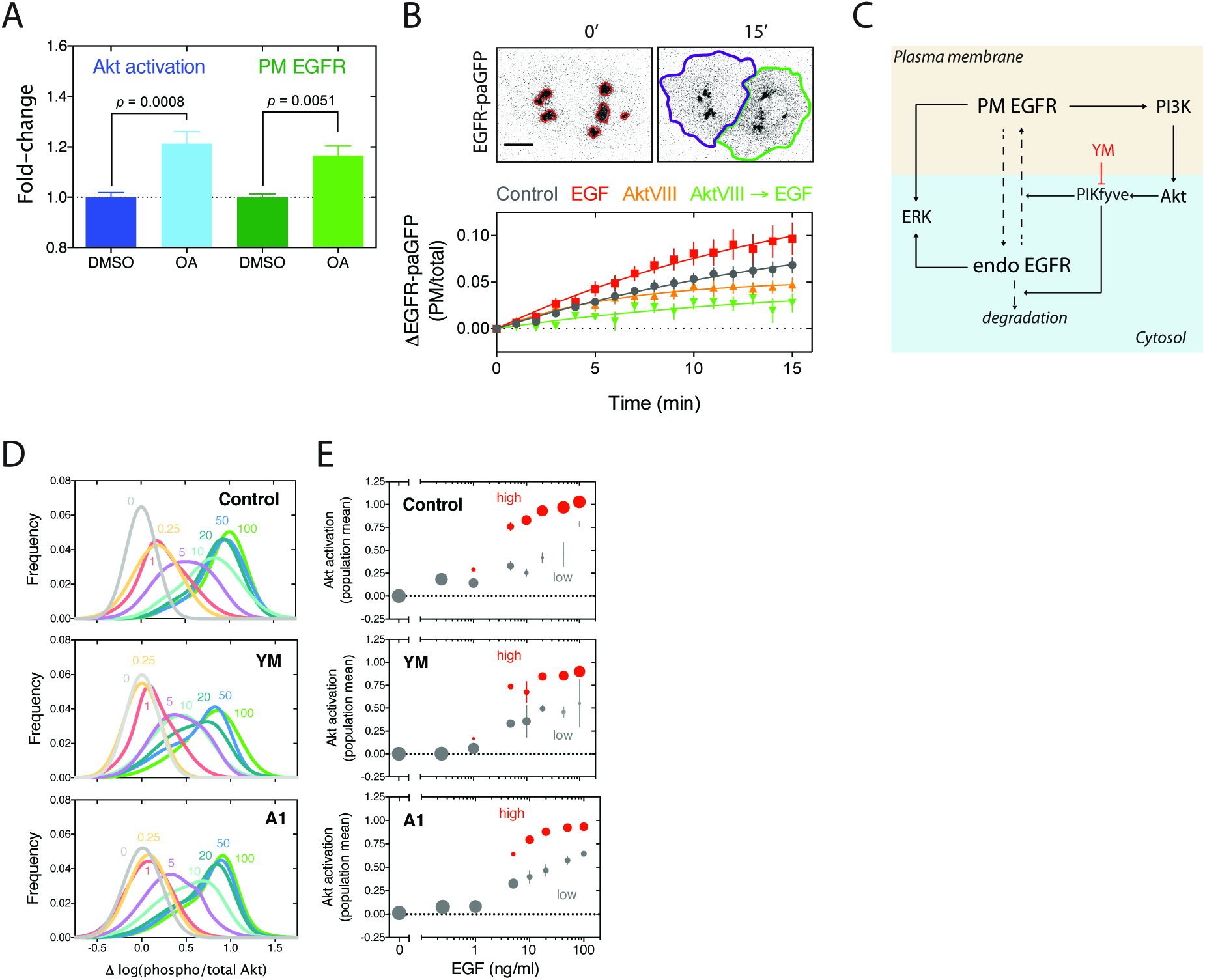
Coupling EGFR activity to vesicular recycling generates a positive feedback. **(A)** Quantification of endogenous PM EGFR abundance and Akt activation in Cos-7 cells by OCW and ICW, respectively, following treatment with the PP2A inhibitor okadaic acid (OA, 1 μM, 2 h) (means ± s.e.m.) **(B)** Representative images and quantification of EGFR-paGFP recycling to the PM following endosomal photoactivation in Cos-7 cells (top) in control, EGF (20 ng/ml, 15 min), AktVIII (10 μM, 1 h) and AktVIII-pretreated, EGF-stimulated (bottom, *N* = 6-10 cells/condition) (means ± s.e.m.) Scale bar = 20 **u**m **(C)** Spatial network topology showing positive feedback generated by coupling PM EGFR activity and Akt-dependent vesicular recycling. PIKfyve inhibition by YM201636 (YM) decouples Akt activation from its effect on EGFR recycling. **(D-E)** Single cell measurements of Akt phosphorylation by flow cytometry in control (top), YM-(200 nM, 1h, middle) and ephrinA1-Fc-(A1, 2 μg/ml, 1h, bottom) pretreated Cos-7 cells endogenously expressing the receptors following stimulation with EGF concentrations indicated (ng/ml, 1 h). (D) Solid lines represent the sum of two Gaussian fits for data accumulated from at least 10 000 cells per condition in 3-4 independent experiments. (E) Quantification of Akt activation for increasing concentrations of EGF. Circle sizes represent the relative proportions of the low (gray) and high (red) Akt activity populations estimated from the Gaussian distributions derived for each EGF concentration shown in D (means ± s.e.m.).

Positive feedback in combination with inhibitory network motifs can convert graded inputs into switch-like, ultrasensitive signaling responses (*44*). Since Akt is preferentially activated at the PM (**Fig. S4**), the EGF-induced increase in EGFR vesicular recycling (**Fig. 4B**) could generate a positive feedback for Akt activation (**Fig. 4C**). To investigate if this positive feedback can generate a switch-like activation of Akt, we measured Akt phosphorylation in thousands of individual Cos-7 cells by flow cytometry following sustained stimulation with a range of EGF concentrations (**Fig. 4D-E**). Cells were stimulated in suspension to negate in situ cell-cell contact as an extrinsic source of variability in Akt activation (**Fig. 3E**). At concentrations ≥ 1 ng/ml, EGF stimulation produced a switch-like activation to a high Akt phosphorylation state in a subpopulation of cells, whose proportion increased with EGF concentration (**Fig. 4D-E top**). Decoupling EGFR activation from its effect on vesicular recycling by PIKfyve inhibition (**Fig. 4D-E** **middle**) or A1 pretreatment (**Fig. 4D-E** **bottom**) did not result in a global decrease in cellular Akt activation but rather reduced the proportion of cells generating a high Akt phosphorylation state, consistent with the inhibition of a positive feedback that produces this switch-like response. Intrinsic cell-to-cell variability in the EGF threshold required to stimulate Akt-dependent vesicular recycling, therefore, determines the proportion of cells that transition to a high Akt activity state at a given EGF concentration. Eph receptor activation, by decoupling EGFR activation from its effect on vesicular trafficking, reduces Akt activation within the population by decreasing the proportion of cells transitioning to a high Akt activity state during EGF stimulation.

#### Eph activation at cell-cell contact suppresses the EGF-promoted transition to a migratory state

EGFR signaling to effectors at the PM generates exploratory cellular behaviors (*45*-*52*) that must be maintained to induce a persistent migratory response. Given that contact inhibitory Eph receptor activation selectively suppresses PM signaling during sustained, subsaturating EGF stimulation (**Fig. 3A**, **4D-E**), we investigated if cell-cell contact regulates EGF-promoted migration by inhibiting Akt-dependent recycling. Since Cos-7 cells exhibit limited migratory behavior, we examined NIH 3T3 mouse embryonic fibroblast (MEF) cells, which generate a haptotactic migratory response to fibronectin that is enhanced by EGF through an increase in exploratory behavior (*53*). Similar to Cos-7 cells (**Fig. 1A-B**), these cells also exhibit a Eph-activity dependent depletion of PM EGFR abundance (**Fig. 5A-B**). Following stimulation with a subsaturating EGF concentration (20 ng/ml), we observed a significant increase in the proportion of migratory cells (**Fig. 5C** **top, Movie S4**, **Fig. S5**), but no change in the average distance travelled per cell (**Fig. 5C** **bottom**). This indicates that EGF promotes the transition of individual cells to a migratory state rather than increasing overall cellular motility. Since EGF binding promotes receptor ubiquitination and degradation, sustained stimulation with supraphysiological saturating EGF concentrations (100 ng/ml) induces a rapid loss in EGF sensitivity with time and thus did not significantly increase the proportion of migratory cells (**Fig. 5C** **top**). Decoupling EGFR activation from its effect on Akt-dependent recycling through the inhibition of PIKfyve or following Eph receptor activation decreased the proportion of migratory cells (**Fig. 5C** **top**). We observed further that increasing concentrations of A1 progressively decreased EGF-induced migration (**Fig. 5C** **top**), consistent with its concentration-dependent effect on EGF-promoted Akt activation (**Fig. 3C**) and suggesting that the amount of ephrinA1-Eph receptor interactions at points of cell-cell contact may determine whether a cell initiates a migratory response to EGF. Indeed, we found that the number of migratory cells following EGF stimulation was inversely proportional to cell density (**Fig. 5D**) and that the increase in migration observed at low densities could be countered by treatment with soluble A1 to mimic Eph receptor contact inhibitory signaling (**Fig. 5D**). Thus, physiological Eph receptor activation at points of homotypic cell-cell contact suppresses EGF-promoted migration by inhibiting Akt-dependent vesicular recycling. However, by preserving endosomal ERK activation following EGF stimulation (**Fig. 3A** **bottom**), we found that neither PIKfyve inhibition nor A1 pretreatment led to a reduction in EGF-promoted cell proliferation (**Fig. 5E**). Thus, by altering the spatiotemporal distribution of EGFR activity, contact inhibitory signaling by Eph receptors influences the cellular outcome to EGF stimulation, preserving a proliferative response while suppressing cell migration.

**Figure 5:**
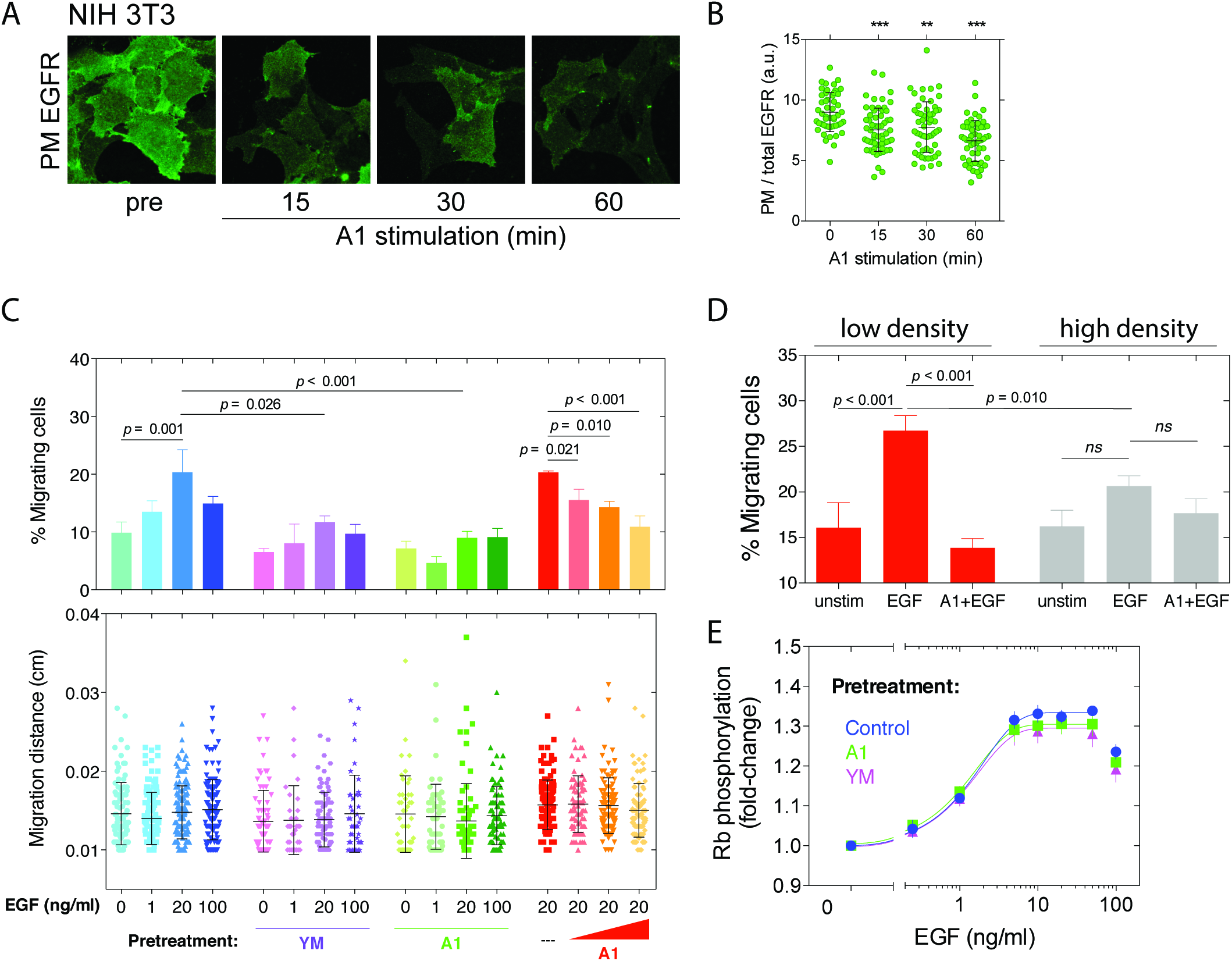
Eph activation at cell-cell contact suppresses the EGF-promoted transition to a migratory state. **(A)** Representative immunofluorescence images and **(B)** quantification of endogenous PM EGFR abundance in NIH 3T3 cells following ephrinA1-Fc (A1, 2 μg/ml) stimulation (means ± sd). Statistical significance was determined using a one-way AN OVA with Sidak’s *post-hoc* test (***, *p <* 0.001; **, *p <* 0.01). **(C)** Percent of NIH 3T3 cells initiating a migratory response (top, means ± s.e.m) and the distance travelled by migratory cells (bottom, means ± s.d.) following EGF stimulation. Cells were pretreated with vehicle (control), YM201636 (YM, 200 nM, 1 h), or ephrinA1-Fc (A1, 2 μg/ml, 1h, green, or 0.02-2 μg/ml as indicated, red) followed by EGF stimulation (0-100 ng/ml as indicated) for 16 h. **(D)** Percent of migrating NIH 3T3 cells when seeded at low or high density in a single well following pretreatment with vehicle or A1 (2 μg/ml, 1 h) and stimulated with EGF (20 ng/ml) for 16 h (means ± s.e.m). Data in C and D were obtained from least three independent experiments, consisting of at least two replicates per experiment (*N* = 581-1483 cells/condition) and statistical significance was determined using an ordinary one-way ANOVA with Holm-Sidak’s multiple corrections *post-hoc* test. **(E)** Quantification of retinoblastoma (Rb) phosphorylation by ICW for vehicle-(control), A1-(2 μg/ml, 1 h) and YM-(200 nM, 1 h) pretreated NIH 3T3 cells following 24 h EGF stimulation at the concentrations indicated (means ± s.e.m).

#### Modulation of vesicular dynamics as a general mechanism to produce context-dependent EGFR signaling

To determine whether environmental signals other than cell-cell contact can influence EGFR signaling through changes in its vesicular trafficking, we investigated the effect of activation of the G protein-coupled receptor Kiss1 (Kiss1R), which, similar to Eph receptors, inhibits Akt (*54*) and suppresses cell migration and metastatic invasion (*55*). Stimulation with the soluble Kiss1R ligand kisspeptin-10 (Kp-10) reduced Akt activity in HEK293 cells and decreased PM EGFR abundance (**Fig. 6A**). Similar to the effect of cell-cell contact, pretreatment with Kp-10 selectively inhibited EGF-promoted Akt activation (**Fig. 6B**), while preserving ERK activation (**Fig. 6C**). The modulation of EGFR vesicular trafficking dynamics could therefore provide a general mechanism to generate plasticity in the signaling response to EGFR activation, through which diverse environmental signals such as cell-cell contact or soluble stimuli like Kp-10 can influence the cellular response to EGF.

**Figure 6:**
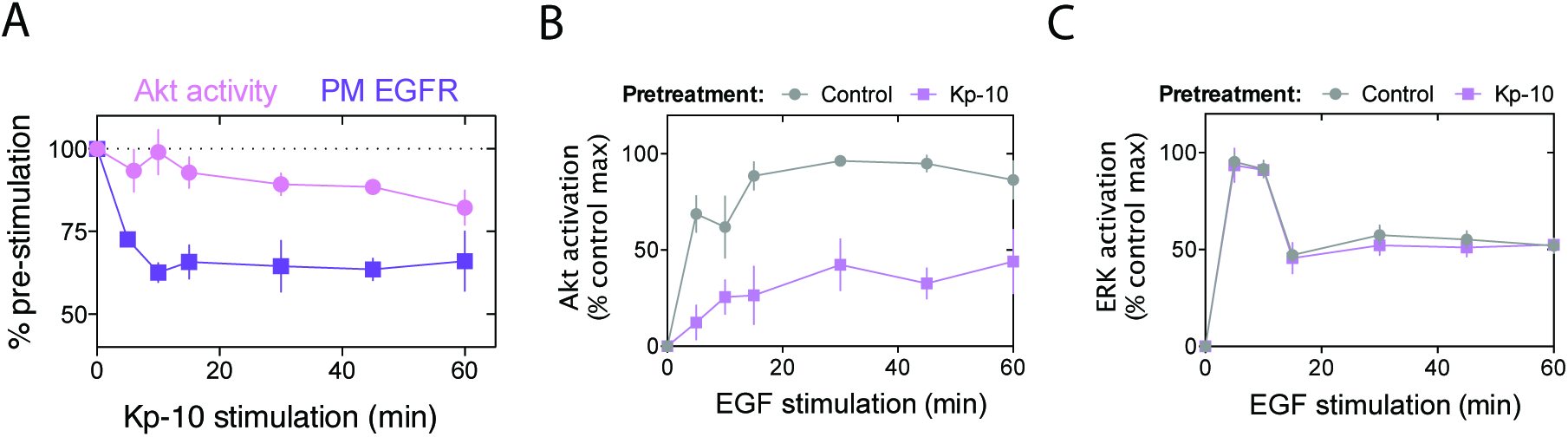
Modulation of vesicular dynamics as a general mechanism to produce context-dependent EGFR signaling. **(A)** Quantification of Akt activation and PM EGFR abundance in HEK293 cells by ICW and OCW, respectively, following stimulation with kisspeptin-10 (Kp-10,100 nM) (means ± s.e.m). **(B,C)** Quantification of Akt and ERK activation by ICW in HEK293 cells for control and Kp-10-pretreated (100 nM, 1 h) cells following EGF stimulation (1 ng/ml) (means ± s.e.m).

## DISCUSSION

In this paper, we demonstrate that Eph receptor activation at cell-cell contacts can generate context-dependent cellular responses to EGF stimulation by modulating EGFR vesicular trafficking dynamics.

Chemotaxis requires that cells remain responsive to stimuli for prolonged periods of time as they migrate toward the chemotactic source. Through an increase in Akt-dependent recycling (**Fig. 4B**), EGF stimulation maintains EGFR density at the PM during persistent, subsaturating stimulation. Since Akt itself is preferentially activated at the PM (**Fig. S3**), the EGF-promoted increase in vesicular recycling generates a positive feedback that switches cells to a high Akt activation state (**Fig. 4D-E**). Although Akt has been previously observed on endosomal membranes through interactions with the early endocytic adaptor protein APPL1 (*56*, *57*), de novo activation of Akt by EGFR requires the production of PI(3,4,5)P_3_, which is impeded by the low abundance of PI(4,5)P_2_ in endosomal membranes (*41*, *58*). Although Akt activation may occur to some extent on endosomal membranes (*59*), since the coupling of active EGFR to Akt activation will be more efficient at the PM, any perturbations that influence the spatial distribution of EGFR, such as Eph receptor activation, will influence the capacity of EGFR to activate Akt (**Fig. 3A**, **Fig. 6A-B**, **Fig. S3**).

We observed that the switch to a high Akt activity state only occurs in a proportion of cells, even in the absence of in situ cell-cell contacts, and increases with EGF concentration (**Fig. 4D-E**). Population heterogeneity in Akt activation has been previously attributed to cell-to-cell variation in PI3K expression (*60*). Our data suggest that intrinsic variability in the expression of signaling and/or trafficking effectors may determine, for a given cell, the EGF concentration required to stimulate Akt-dependent trafficking and engage the positive feedback that produces a high Akt activity state. Small differences in EGF concentration substantially influence the proportion of cells generating a high Akt response (e.g. a shift from 5 to 10 ng/ml increases the proportion of cells from 43 to 85%, respectively, **Fig. 4D-E**). Perhaps it is not coincidental that the concentration range over which this switch occurs corresponds to the physiological range of EGF concentrations (*29*). By generating a sharp boundary for Akt activation within the physiological EGF concentration regime, even slight changes in the threshold of this switch could have profound implications for tissue dynamics (e.g. initiation of migration).

Eph receptor activation, for example, by suppressing EGFR recycling, decreased the proportion of cells generating a high Akt response from 85 to 41% in response to 10 ng/ml EGF (**Fig. 4D-E**). The dependence of Akt activation on EGFR recycling allows the degree of cell-cell contact to regulate the proportion of cells generating a migratory response to EGF stimulation. PI3K/Akt signaling has previously been suggested as the point of convergence for EGFR/Eph control of cell migration (*15*); however, the molecular mechanism underlying this oppositional relationship remained unclear. Our results indicate that Eph receptor activation inhibits EGF-promoted cell migration by suppressing Akt-dependent recycling, thus impeding the spatially-maintained positive feedback that generates a high Akt response and decreasing the sensitivity of cells to persistent EGF stimulation necessary to maintain exploratory behavior. However, by changing the spatial distribution of EGFR activity (**Fig. 2C-D**), Eph receptor activation selectively suppresses migratory signaling from the PM while leaving proliferative ERK signaling intact (**Fig. 3A**, **Fig. 3E-F**, **Fig. 5C-D**). This contextual plasticity generates two distinct cellular outcomes to EGF stimulation that may be important in physiological settings such as wound healing. At the tissue boundary, cells with reduced cell-cell contact would increase their exploratory behavior in response to EGF released at the site of the wound. Cells located deeper in the tissue, despite extensive cell-cell contact, would retain their proliferative response to extracellular EGF, and undergo mitosis to fill the vacant space created as exploratory cells migrate to occupy the wound area.

Our observations demonstrate that communication between receptors with opposed functionality can emerge through changes in vesicular trafficking dynamics rather than relying on direct interactions between the receptors or their respective effectors. Such a mechanism also allows different receptors with similar functional roles (e.g. EphA2 and Kiss1R) to alter the cellular response to stimuli without having to evolve distinct protein interaction domains to do so. The dependency of EGFR signaling on its vesicular dynamics could confer a general mechanism through which the cell can generate functional plasticity to growth factor stimulation while preserving specificity in cell-cell communication.

## Materials and methods

### Primary antibodies

Mouse anti-Akt (2920, Cell Signaling Technology (CST), Danvers, MA, USA), mouse anti-Akt-Alexa488 (2917, CST), rabbit anti-phospho-Akt(Ser^473^) (4060, CST), rabbit anti-phospho-Akt (Ser^473^)-Alexa647 (4075, CST), mouse anti-HA (9658, Sigma-Aldrich, St.Louis, MO, USA), rabbit anti-EGFR (4267, CST), goat anti-EGFR (AF231, R&D Systems, Minneapolis, MN, USA), mouse anti-phospho-EGFR(Tyr^845^) (558381, BD Biosciences, Heidelberg, Germany), rabbit anti-phospho-EGFR(Tyr^1045^) (2237, CST), mouse anti-phospho-EGFR(Tyr^1068^) (2236, CST), goat anti-EphA2 (R&D Systems), rabbit anti-phospho-Eph(Tyr^588/596^) (Abcam, Cambridge, UK), mouse anti-ERK1/2 (4696, CST), rabbit anti-phospho-ERK(Thr^202/Tyr204^) (4370, CST), mouse anti-Rab5 (610724, BD Biosciences), rabbit anti-Rab7 (9367, CST), rabbit anti-phospho-Rb(Ser807/811, CST) mouse anti-tubulin (6074, Sigma-Aldrich)

### Secondary antibodies

IRDye 680RD Donkey anti-Mouse (LI-COR Biosciences), IRDye 680RD Donkey anti-Rabbit (LI-COR Biosciences), IRDye 680RD Donkey anti-Goat (LI-COR Biosciences), IRDye 800CW, Donkey anti-Mouse (LI-COR Biosciences), IRDye 800CW Donkey anti-Rabbit (LI-COR Biosciences), IRDye 800CW Donkey anti-Rabbit (LI-COR Biosciences), AlexaFluor 405 goat anti-Mouse (Life Technologies), AlexaFluor 488 donkey anti-Goat (Life Technologies), AlexaFluor 546 donkey anti-rabbit (Life Technologies), AlexaFluor 647 donkey anti-Rabbit (Life Technologies)

### Plasmids

Generation of EGFR-mCitrine, EGFR-mCherry, EGFR-paGFP, PTB-mCherry, c-Cbl-BFP and HA-ubiquitin (*25*), as well as EphA2-mCitrine, SH2-mCherry and LIFEA2 (*24*) were previously described. pcDNA3.1-EphA2 was a gift from Tony Pawson.

### Reagents

AktVIII (sc-3513, Santa Cruz Biotechnology, Dallas, TX, USA), EGF (AF-100-15, Peprotech, Hamburg, Germany), okadaic acid (sc-3513, Santa Cruz Biotechnology), YM201636 (13576, Biomol GmbH, Hamburg, Germany), dynole 31-2 (ab120464, Abcam), dynole 34-2 (ab120463, Abcam), Kisspeptin-10 (445888, Merck Millipore). EGF-Alexa647 was prepared as previously described (*25*). EphrinA1-Fc (602-A1-200) was preclustered by incubating with chicken Anti-Fc (GW200083F, Sigma-Aldrich) at a ratio of 5:1 at room temperature for at least 30 min.

### Cell culture

Cos-7, HEK293 and NIH 3T3 cells were grown in Dulbecco’s Modified Eagle’s Medium (DMEM) supplemented with 10% fetal bovine serum (FBS), 2 mM L-glutamine and 1% non-essential amino acids (NEAA) and maintained at 37**°**C in 5% CO_2_. MCF10A cells were grown in DMEM/F12 supplemented with 5% horse serum, 20 ng/ml EGF, 500 ng/ml hydrocortisone, 100 ng/ml cholera toxin and 10 μg/ml insulin and maintained at 37**°**C in 5% CO_2_. MDA-MB-231 cells were grown in Leibowitz medium supplemented with 10% FBS and 2mM L-glutamine maintained at 37**°**C in 0% CO_2_. When required, transfection of cells was performed using FUGENE6 (Roche Diagnostics, Mannheim, Germany) or Lipofectamine 2000 (Life Technologies, Darmstadt, Germany) according to manufacturer’s protocol. Approximately 16-18 hours prior to an experiment, cells were starved in DMEM containing 0.5% FBS, 2 mM L-glutamine and 1% NEAA. One hour before stimulation, starvation media was changed to serum-free DMEM or DMEM without phenol red for live cell imaging. Unless explicitly stated in the figure legends, all experiments were performed with Cos-7 cells with endogenous expression of EGFR and Eph receptors.

### In-cell westerns (ICW) and on-cell westerns (OCW)

Cells were seeded on black, transparent bottomed 96-well plates (3340, Corning, Hagen, Germany) coated with poly-L-lysine (P6282, Sigma Aldrich). Cell were fixed with Roti-Histofix 4% (Carl Roth, Karsruhe, Germany) for 5 min at 37°C. For ICW, cells were permeabilized with 0.1% Triton X-100 (v/v) for 5 min at room temperature. For OCW, cells were not permeabilized. Samples were incubated in Odyssey TBS blocking buffer (LI-COR Biosciences, Lincoln, NE, USA) for 30 min at room temperature. Primary antibodies were incubated overnight at 4°C and secondary antibodies (IRDyes, LI-COR Biosciences) were incubated in the dark for 1 h at room temperature. All wash steps were performed with TBS (pH 7.4). Intensity measurements were made using the Odyssey Infrared Imaging System (LI-COR Biosciences). ICW/OCW were calibrated by Western blots to ensure accurate quantification. Quantification of the integrated intensity in each well was performed using the MicroArray Profile plugin (OptiNav Inc., Bellevue, WA, USA) for ImageJ v1.47 (http://rsbweb.nih.gov/ij/). In each ICW or OCW, 2-4 replicates per conditions were obtained per experiment, and all data presented represents means ± s.e.m. from at least three independent experiments.

### Immunofluorescence

Cells were cultured on 4-or 8-well chambered glass slides (Lab-tek, Thermo Fisher Scientific, Waltham, MA) and fixed with 4% paraformaldehyde/PBS (w/v) for 10 min at 4°C. To measure PM EGFR, fixed, non-permeabilized samples were first incubated with primary antibody directed at an extracellular epitope of EGFR (AF231, R&D Systems, 1:200) overnight at 4°C followed by secondary antibody for 1 h at room temperature. For all other immunofluorescence experiments, samples were permeabilized with 0.1% Triton X-100 (v/v) for 5 min at room temperature prior to incubation with primary antibodies. All wash steps were performed with TBS (pH 7.4). Fixed samples were imaged in PBS at 37°C. For all analysis, an initial background subtraction was performed on immunofluorescence images. To quantify the proportion of EGFR in Rab5 and Rab7 compartments, binary masks were generated from intensity thresholded images of Rab5 and Rab7 staining. To generate a mask of Rab5/Rab7 double positive endosomes, the product of their individual masks was used. The integrated fluorescence intensity of EGFR-mCherry was determined in each of the endosomal masks and divided by the total integrated EGFR fluorescence intensity of the cell. All analysis was performed using ImageJ. A cell segmentor tool was developed in-house in Anaconda Python (Python Software Foundation, version 2.7, https://www.python.org/) to quantify the spatial distribution of EGFR and pTyr845-EGFR in fixed cells. Cells were divided into 6 equally spaced radial bins emanating from the center of cell mass.

### Confocal imaging

Routinely, cells were cultured for live cell confocal imaging on 4-or 8-well chambered glass slides (Lab-tek) and transiently transfected as described above. Confocal images were recorded using an Olympus Fluoview FV1000 confocal microscope (Olympus Life Science Europa, Hamburg, Germany) or a Leica SP8 confocal microscope (Leica Microsystems, Wetzlar, Germany).

### Olympus Fluoview™ FV1000

The Olympus Fluoview™ FV1000 confocal microscope was equipped with a temperature controlled CO2 incubation chamber at 37°C (EMBL, Heidelberg, Germany) and a 60x/1.35 NA Oil UPLSApo objective (Olympus, Hamburg, Germany). EphA2-mCitrine and EGFR-mCherry were excited using a 488 nm Argon-laser (GLG 3135, Showa Optronics, Tokyo, Japan) and a 561 nm DPSS laser (85-YCA-020-230, Melles Griot, Bensheim, Germany), respectively. Detection of fluorescence emission was restricted with an Acousto-Optical Beam Splitter (AOBS) for mCitrine @ 498-551 nm and mCherry @ 575-675 nm. In all cases, scanning was performed in frame-by-frame sequential mode with 2x frame averaging. The pinhole was set to 250 μm.

### Leica SP8

The Leica TCS SP8 confocal microscope was equipped with an environment-controlled chamber (LIFE IMAGING SERVICES, Switzerland) maintained at 37°C and a HC PL APO CS2 1.4 NA oil objective (Leica Microsystems, Wetzlar, Germany). Alexa488-conjugated secondary antibodies, fluorescent fusion proteins containing mCitrine and mCherry, and EGF-Alexa647 were excited using a 470-670 nm white light laser (white light laser Kit WLL2, NKT Photonics, Denmark) at 488, 514 and 561 and 647 nm, respectively. PH-Akt-Cerulean was excited using an Argon Laser at 458 nm. Detection of fluorescence emission was restricted with an AOBS as follows: Cerulean @ 468-505 nm, Alexa488 @ 498-551 nm, mCitrine @ 525-570 nm, mCherry @ 570-650 nm and Alexa647 @ 654-754 nm. The pinhole was set to 250 μm and 12-bit images of 512x512 pixels were acquired in a frame-by-frame sequential mode.

### Analysis of time-lapse confocal imaging

All analysis of live cell imaging data required an initial background subtraction for all images obtained.

To quantify the proportion of endosomal EGFR-mCherry or EphA2-mCitrine, binary masks of endosomes were generated from intensity thresholded images. The integrated fluorescence intensity of EGFR-mCherry and EphA2-mCitrine was determined in their corresponding endosomal masks and divided by the total integrated fluorescence intensity of the cell.

Fluorescence localization after photoactivation (FLAP) experiments were carried out at 37°C on a Leica SP8. EGFR-mCherry was co-expressed to identify and select regions of endosomal EGFR for photoactivation. Background intensity of EGFR-paGFP prior to photoactivation was measured and subtracted from post-activation images. Photoactivation of EGFR-paGFP was performed with the 405nm laser at 90% power. Following photoactivation, fluorescence images of EGFR-paGFP were acquired every minute for a total of 15 minutes. PM EGFR-paGFP fluorescence was quantified as the integrated intensity in a 5-pixel ring of the cell periphery and, after subtracting pre-activation background intensity, was calculated as a proportion of total EGFR-paGFP intensity.

### Fluorescence lifetime imaging microscopy (FLIM)

EGFR-mCitrine, PTB-mCherry and HA-c-Cbl-BFP were ectopically expressed in Cos-7 cells. Fluorescence lifetime measurements of EGFR-mCitrine were performed at 37°C on a Leica SP8 equipped with a time-correlated single-photon counting module (LSM Upgrade Kit, Picoquant, Berlin, Germany) using a 63x/1.4 NA oil objective. EGFR-mCitrine was excited using a pulsed WLL at a frequency of 20 MHz and fluorescence emission was restricted with an AOBS to 525-570 nm. Photons were integrated for a total of ~ 2 min per image using the SymPhoTime software V5.13 (Picoquant, Berlin, Germany). Data analysis was performed using custom software in Anaconda Python based on global analysis as described in (*61*).

Fluorescence lifetime measurements of LIFEA2 were performed and analyzed as previously described (*24*).

### Flow cytometry

Cells were detached using accutase, centrifuged at 200g for 5 min and resuspended in serum-free DMEM. Cells were fixed with 5% sucrose/Roti-Histofix (w/v) for 15 min at 37°C. Ice-cold methanol was added to 90% (v/v) for 30 min on ice. Cells were rinsed once with 0.5% BSA/TBS (w/v) and incubated with Odyssey TBS blocking buffer (LI-COR Biosciences) for 30 min at room temperature. Anti-phospho-Akt(Ser^473^)-Alexa647 (4075, Cell Signaling Technology) was added directly to blocking buffer and incubated overnight at 4°C. Anti-Akt-Alexa488 (2917, Cell Signaling Technology) was added for 2 h prior to measurement. Samples were analyzed using the LSRII flow cytometer (BD Biosciences). Alexa488 was excited with a 488nm laser and fluorescence emission was collected using a 505 nm LP dichroic and a 530/30 nm filter. Alexa647 was excited with 633 nm lasers and fluorescence emission was collected using a 670/40 nm filter. Samples were analyzed using FlowJo v10 (FlowJo, LLC, Ashland, OR, USA) to obtain single cell intensity measurements of phospho-and total Akt. Population distributions of log(phospho/total Akt) were fitted with a single Gaussian or a sum of two Gaussian distributions using GraphPad Prism (GraphPad Software, La Jolla, CA, USA).

### Cell migration

NIH 3T3 cells were seeded onto fibronectin-coated (F0895, Sigma, 1.25 ug/cm^2^c) 12 well culture dishes (83.3921, Sarstedt, Nuembrecht, Germany) containing 2-well Culture-Inserts (80209, ibidi) to create a cell-free area. Immediately before stimulation, inserts were removed and cells were incubated with Hoechst to label nuclei. Wide field images were acquired using an Olympus IX81 inverted microscope (Olympus, Hamburg, Germany) equipped with a MT20 illumination system, a 4x/0.16 NA air objective and an Orca CCD camera (Hamamatsu Photonics, Hamamatsu City, Japan). Transmission and fluorescence images were acquired every 10 min for 16 h. The cell-free area created by the Culture-Insert was cropped using ImageJ and defined as the migration region. Individual cells were detected and tracked by their nuclear Hoechst staining as they travelled within the migration region using the TrackMate ImageJ plugin (*62*), and the total distance of each track was quantified.

### Immunoprecipitation (IP) and western blotting

Cells were lysed in TGH (150 mM NaCl, 2 mM EGTA/EDTA, 50 mM HEPES (pH 7.4), 1% Triton X-100,10% glycerol, 1 mM phenylmethylsulfonyl fluoride, 10 mM N-ethylmaleimide(NEM)) or RIPA (for immunoprecipitation; 50 mM Tris-HCl (pH 7.5), 150 mM NaCl, 1 mM EGTA, 1 mM EDTA, 1% Triton X-100,1% Sodium deoxycholate, 0.2% SDS, 2.5 mM sodium pyrophosphate and 10 mM NEM), supplemented with Complete Mini EDTA-free protease inhibitor (Roche Applied Science, Heidelberg, Germany) and 100 μl phosphatase inhibitor cocktail 2 and 3 (P5726 and P0044, Sigma Aldrich). Lysates were sonicated prior to centrifugation at 14 000 rpm for 10 min at 4°C to separate non-soluble material. For immunoprecipitation, cell lysates were incubated with 50 μl washed Protein G magnetic beads (10003D, Life Technologies) for 1 h at 4°C to pre-clear the samples from unspecific binding proteins. Supernatants were incubated with primary antibody alone for 2 h followed by the addition and overnight incubation with Protein G magnetic beads at 4°C with agitation. SDS-PAGE was performed using the X-cell II mini electrophoresis apparatus (Life Technologies) according to the manufacturer’s instructions. Samples were transferred to preactivated PVDF membranes (Merck Millipore, Billerica, MA) and incubated with the respective primary antibodies at 4**°**C overnight. Detection was performed using species-specific secondary IR-Dye secondary antibodies (LI-COR Biosciences) and the Odyssey Infrared Imaging System (LI-COR Biosciences). The integrated intensity of protein bands of interest was measured using the ImageJ software and signals were normalised by dividing the intensities of phosphorylated protein by total protein intensities or by dividing intensities of co-immunoprecipitated proteins by the corresponding immunoprecipitated protein.

## Acknowledgments

We would like to thank Dr. Astrid Krämer for critically reading this manuscript, Dr. Aneta Koseska for assistance in data analysis and Dr. Klaus Schuermann for generating the 3-D spatial-temporal maps.

## Funding

The project was partially funded by the European Research Council (ERC AdG 322637) to P.I.H.B.

## Author contributions

P.I.H.B. and W.S. conceived the project. W.S. performed and analyzed most experiments. O.S. performed experiments with LIFEA2 and contributed to the migration experiments, Y.B. contributed to the immunofluorescence experiments, L.B. contributed to the ICW and immunoprecipitation experiments. W.S. wrote the manuscript with assistance from P.I.H.B.

## Competing interests

The authors declare that no competing interests exist.

## Data and material availability

Data, scripts and reagents are available upon request.

## Supplementary Materials

**Figure S1:**
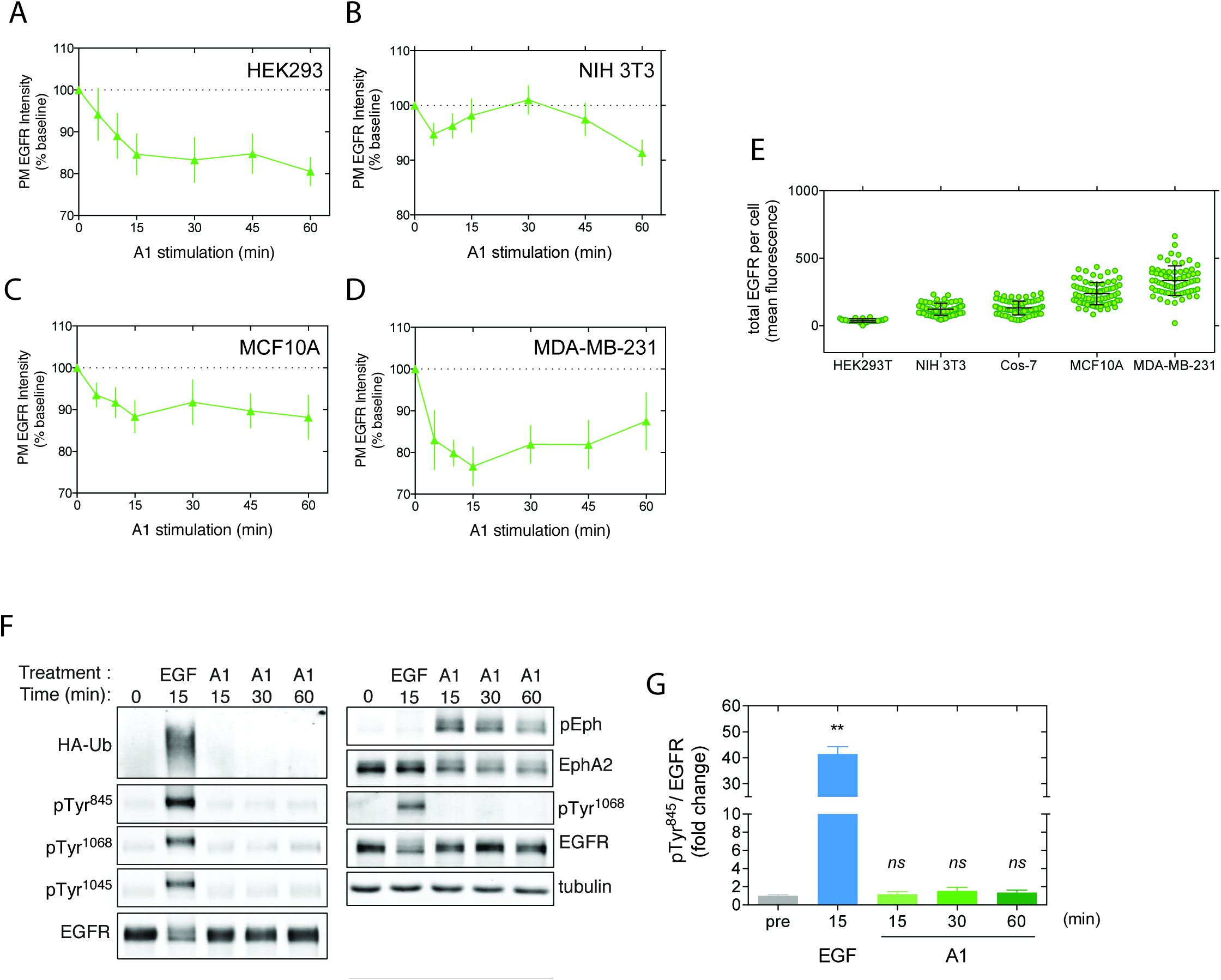
Eph receptor activation reduces PM EGFR abundance. (A-D) Quantification of endogenous PM EGFR abundance by ICW following ephrinA1-Fc stimulation (A1, 2 μg/ml) of (A) HEK293, (B) NIH 3T3, (C) MCF10Aand (D) MDA-MB-231 cells. (means ± s.e.m) (E) Single cell immunofluorescence measurements of EGFR expression in the cell lines used in this study. (F-G) Cos-7 lysates immunoprecipitated (IP) with anti-EGFR (left) or blotted for total proteins (right) following stimulation with EGF (100 ng/ml) or A1 (2 μg/ml) for the indicated times. IP was probed with anti-HA (to detect co-transfected HA-ubiquitin), anti-pTyr^845^, anti-pTyr^1068^, anti-pTyr^1045^ (to detect phosphorylated EGFR) and anti-EGFR. Total lysates were probed with anti phospho-Eph (pEph), anti-EphA2, anti-pTyr^1068^, anti-EGFR and anti-tubulin. Shown are (F) representative blots and (G) quantification of EGFR phosphorylation (pTyr^845^) from four independent experiments (means ± s.e.m). Statistical significance was determined using a one-way ANOVA with Sidak’s *post-hoc* test (**, *p <* 0.01).

**Figure S2:**
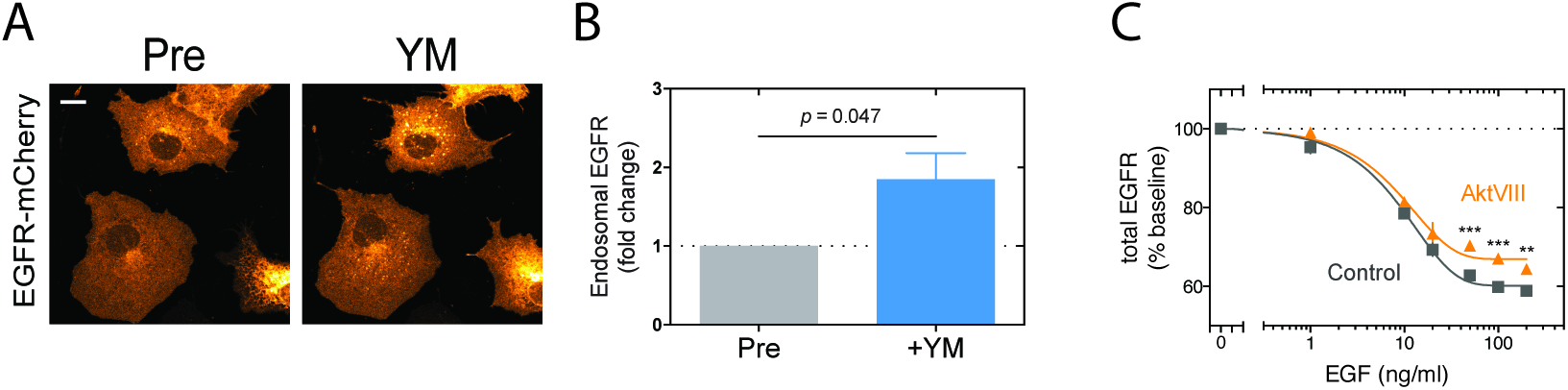
Akt/PIKfyve regulates EGFR vesicular trafficking. (A) Representative images and (B) quantification of endosomal EGFR-mCherry following treatment with the PIKfyve inhibitor YM201636 (YM, 200 nM, 1h) (*N* = 9, mean ± s.e.m). Statistical significance was determined using a two-tailed Student’s t test. Scale bar = 20 μm. (C) ICW measurements of total EGFR abundance following EGF stimulation (1-200 ng/ml, 30 min) in vehicle (control) and AktVIII (10 μM, 1 h) pretreated cells (means ± s.e.m). Statistical significance was determined using a two-way ANOVA with Sidak’s multiple corrections *post-hoc* test (***, *p >* 0.001; **, p > 0.01).

**Figure S3:**
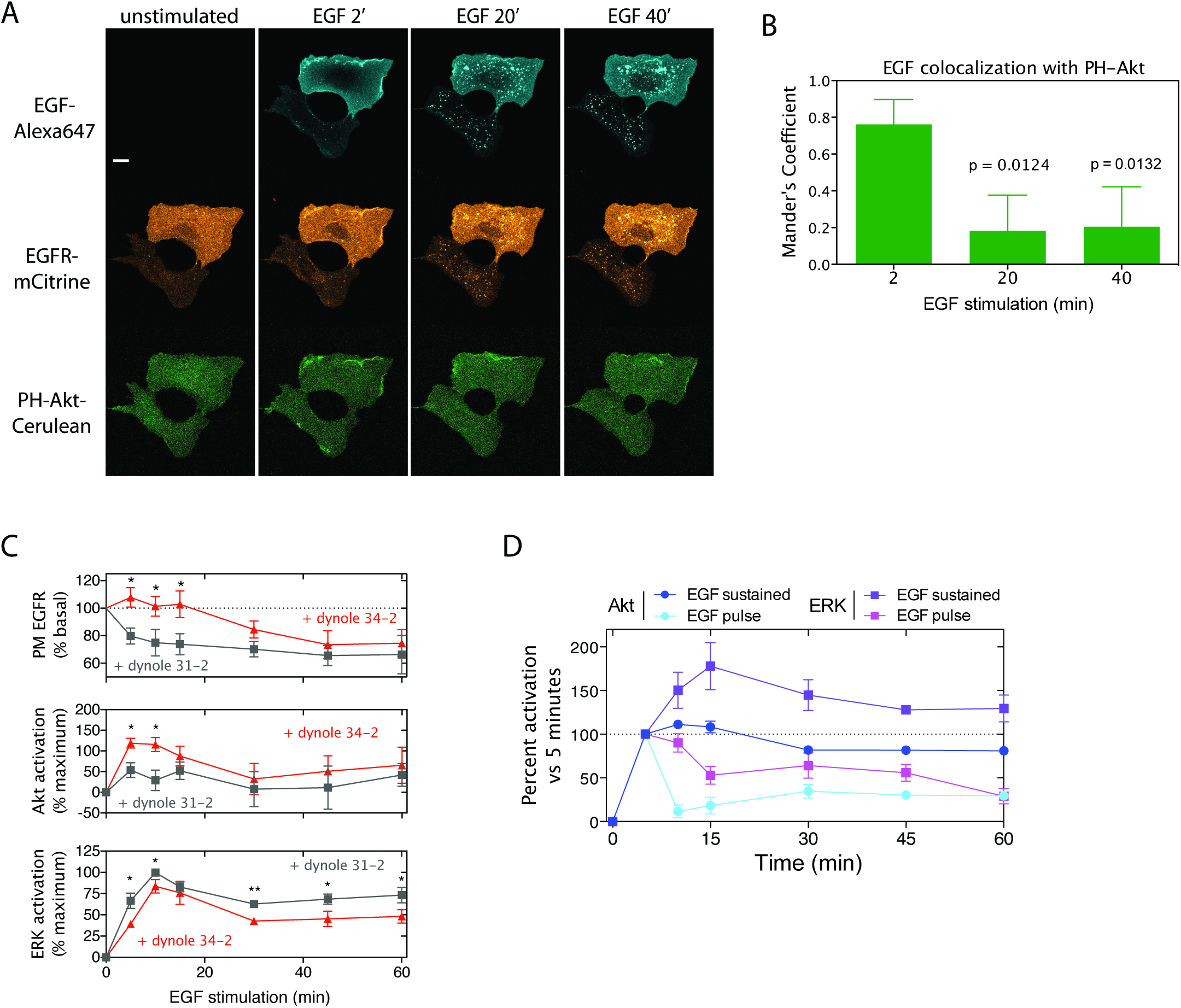
Akt is preferentially activated at the PM following EGFR stimulation. (A)Akt rapidly translocates to regions of EGF-bound receptors at the PM, but is not recruited to endosomes following internalization of ligand-bound receptor. Shown are representative images of Cos-7 cells expressing EGFR-mCitrine and PH-Akt-Cerulean following stimulation with EGF-Alexa647 (100 ng/ml) for the indicated times. (B) Colocalization between EGF-Alexa647 and PH-Akt-Cerulean is highest immediately after stimulation (2 min) while most EGFR-mCitrine is localized at the plasma membrane. As EGF-Alexa647 accumulates in endosomes (20 and 40 min), colocalization with PH-Akt-Cerulean decreased. Quantification of colocalization by Mander’s coefficient from EGF-Alexa647 intensity-dependent thresholded images (means ± sd, *N* = 4). (C) Inhibition of endocytosis increases EGF-promoted Akt activation, while ERK activation is decreased. PM EGFR abundance was quantified by OCW and Akt/ERK activation by ICW in HEK293 cells stimulated with EGF (1 ng/ml) for the indicated times following pretreatment with the dynamin inhibitor dynole 34-2 (100 μM, 30 min) or its negative control analog dynole 31-2 (100 μM, 30 min) (means ± s.e.m). (D) Akt activation is rapidly terminated following EGF washout in Cos-7 cells, indicating the necessity of PM EGF binding for persistent Akt activation. ERK activation decays much slower following EGF removal through due to persistent activity of endosomal EGFR. Akt and ERK activation were measured by ICW following sustained EGF (1 ng/ml) stimulation or a 5 min EGF pulse and subsequent washout (means ± s.e.m). Statistical significance was determined in B using a repeated measures one-way ANOVA with a Dunnett’s *post-hoc* test, and in C using a two-way ANOVA with Sidak’s *post-hoc* test. (***, *p <* 0.001; **, p < 0.01; *, p < 0.05). Scale bars = 20 μm

**Figure S4:**
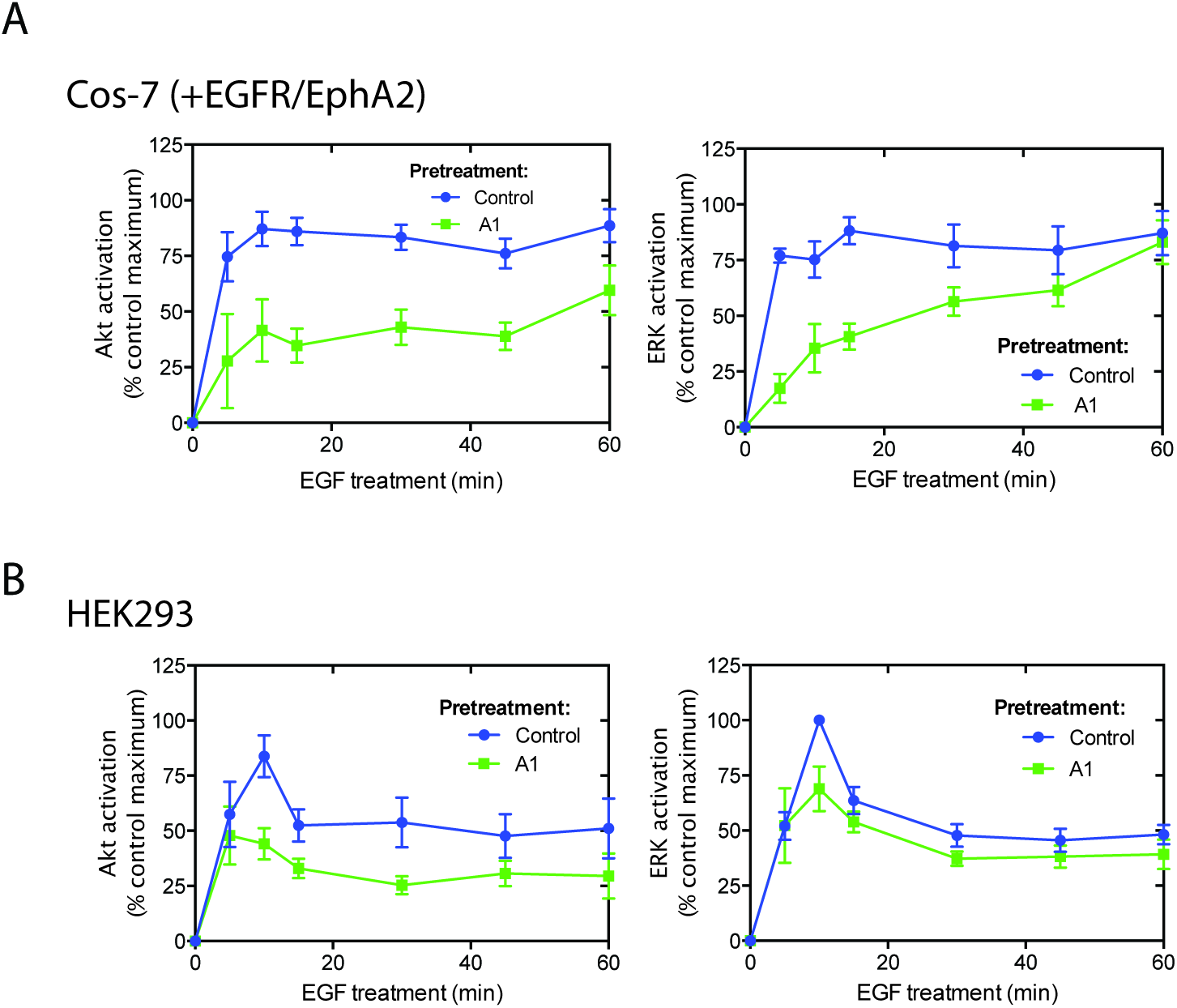
EphrinA1-Fc pretreatment inhibits EGFR-promoted Akt activation. Akt and ERK activation were quantified by ICW in (A) Cos-7 cells ectopically expressing EGFR and EphA2, and in (B) HEK293 cells endogenously expressing the receptors. Cells were pretreated with vehicle (control) or ephrinA1-Fc (A1, 2 μg/ml) for 1 h, followed by 1 ng/ml EGF stimulation for the indicated times. Data represent means ± s.e.m.

**Figure S5:**
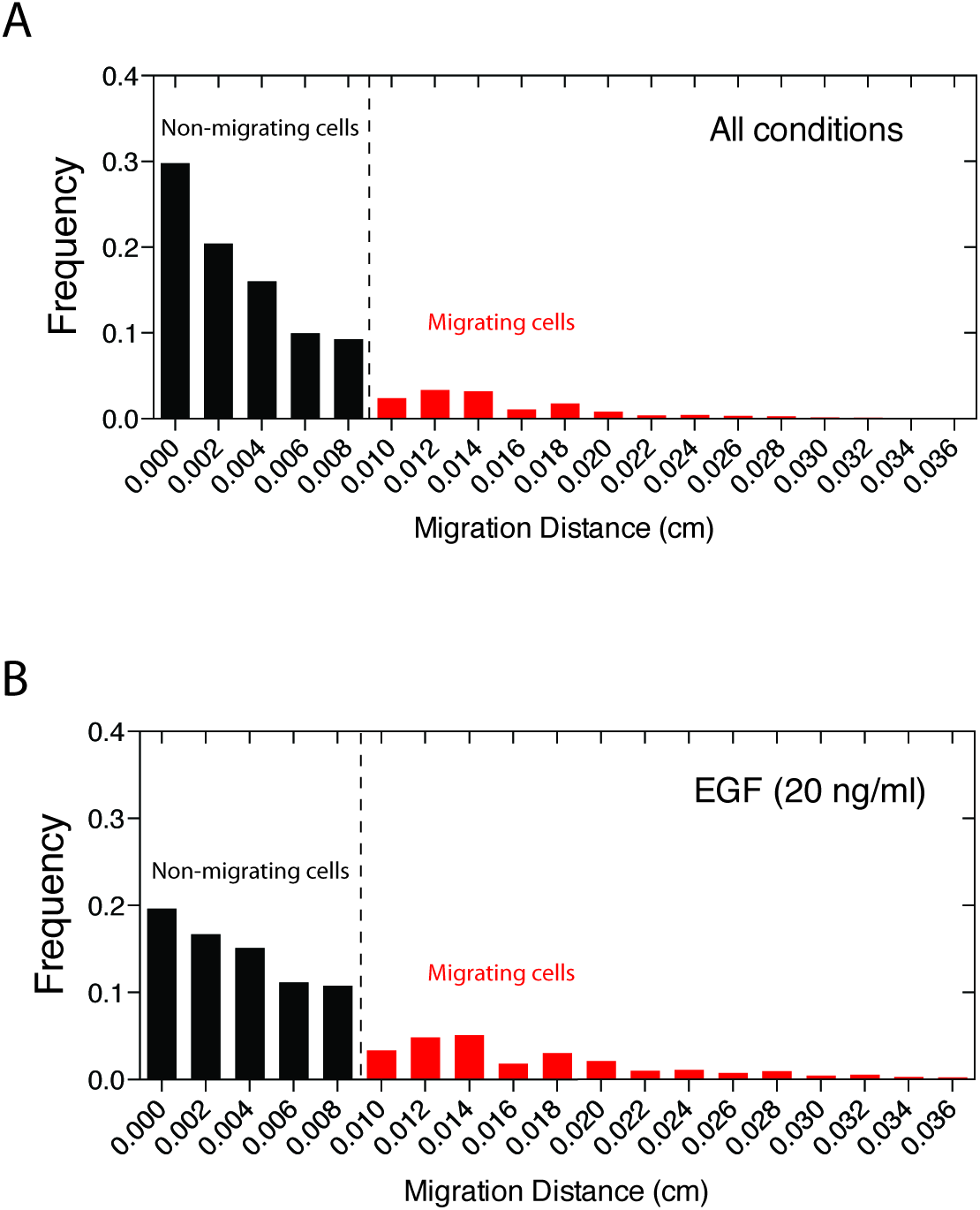
Distributions of cell migration distances. Migration distance was quantified for individual cells as described in *Methods*. Shown are the distributions of migration distance for (A) all tracked cells (*N* = 32 203 cells) and (B) 20 ng/ml EGF treated cells (*N* = 2343 cells). To distinguish migrating cells from those that are pushed into the cell-free area by population expansion due to cell division during the 16 h acquisition time, a minimal displacement distance of 0.01 cm was used as a threshold.

**Movie S1: Akt inhibition induces EGFR endosomal accumulation**. Treatment of Cos-7 cells ectotopically expressing EGFR-mCherry with the Akt inhibitor AktVIII (10 μM) promotes an increase in endosomal EGFR. Scale bar = 20 μm

**Movie S2: EphrinA1-Fc stimulation induces EGFR endosomal accumulation**. Stimulation of Cos-7 cells ectotopically expressing EGFR-mCherry and EphA2-mCitrine with EphrinA1-Fc (2 μg/ml) promotes an increase in endosomal EGFR. Scale bar = 20 μm

**Movie S3: EphrinA1-Fc:EphA2 interactions at cell-cell contact induces EGFR endosomal accumulation**. HEK293 cells ectopically expressing EBFP-EphrinA1 in suspension were added to adherent Cos-7 cells ectopically expressing EGFR-mCherry and EphA2-mCitrine 30 min prior to imaging. Time lapse imaging begins as EBFP-EphrinA1-HEK293T cells make initial cell-cell contact with Cos-7 cells. Scale bar = 20 μm

**Movie S4: EGF-promoted migration in NIH 3T3 cells**. Unstimulated (unstim), EGF (20 ng/ml), YM201636 (200 nM) followed by EGF (20 ng/ml) stimulation (YM) and EphrinA1-Fc (2 μg/ml) followed by EGF (20 ng/ml) stimulation (A1). Scale bar = 100 μm

